# ERM proteins antagonistically control cancer invasion and metastasis via mechanosensing regulation

**DOI:** 10.64898/2026.11.16.767111

**Authors:** Malak Amer, James V. McCann, Tina N. Shogan, Andrew J. Ewald, Haguy Wolfenson

**Author notes:** Correspondence to Haguy Wolfenson. J.V. McCann and T.N. Shogan contributed equally to this work.

## Abstract

Metastasis requires cancer cells to overcome anchorage dependence and rewire mechanosensing pathways. Here, we identify the ERM proteins ezrin, radixin, and moesin as key but antagonistic regulators of breast cancer metastasis. Expression profiling across matched low- and high-metastatic breast cancer models revealed consistent upregulation of all three ERM proteins in highly metastatic cells. Despite this, ezrin depletion reduced colony formation, invasion, and lung metastasis, whereas radixin or moesin depletion enhanced these phenotypes. Loss of radixin or moesin increased ezrin localization at the cell edge, while ezrin depletion reduced membrane-associated moesin, indicating a dynamic balance among ERM proteins. Ezrin depletion also increased focal adhesion size and number, FAK activation, and force transmission to the extracellular matrix, whereas radixin or moesin silencing produced opposite effects. Cellular force magnitude was inversely correlated with invasion, and pharmacological modulation of contractility reverted the effects of ERM depletion. These findings reveal an ERM-dependent mechanosensing checkpoint that regulates metastasis through control of adhesion dynamics and force transmission.

**Summary:** Ezrin, radixin, and moesin form a mechanosensing checkpoint that antagonistically regulates breast cancer metastasis. Ezrin promotes invasion and metastatic colonization, whereas radixin and moesin suppress them by increasing adhesion maturation, cellular force transmission, and ERM redistribution at the cell edge.

## Introduction

Metastasis, the spread and colonization of cancer cells throughout the body, is the primary cause of cancer death (Nicolson, 1979). The detachment of metastasizing cancer cells from the primary tumor, and their subsequent growth in distant organs requires them to be anchorage-independent (Guadamillas et al., 2011; Taddei et al., 2012), which is a critical indicator of aberrant mechanosensing of the extracellular matrix (ECM) (Amer et al., 2021a; Wolfenson et al., 2016a; Yang et al., 2020a). During mechanosensing, cells probe the mechanical properties of the ECM by exerting actomyosin-based forces onto it through integrin-based adhesions. The ensuing mechanical feedback triggers mechanotransduction signaling pathways, thus regulating cellular functions (Geiger & Bershadsky, 2002; Iskratsch et al., 2014; Luo et al., 2013; Sala et al., 2024). Mechanosensing regulatory proteins were shown to be dysregulated in many cancers (Bharadwaj et al., 2005a; Hughes et al., 2003; Pawlak et al., 2004; Yang et al., 2016), causing the cancer cells to apply disproportionate forces and to respond abnormally to the mechanical environment (Amer et al., 2021b; Guadamillas et al., 2011). Restoring their expression levels has been found to re-establish normal mechanosensing activity and revert the cells to an anchorage-dependent state (Bharadwaj et al., 2005b; Stehn et al., 2006; Yang et al., 2020b).

The adhesome is a large network of proteins involved in adhesion regulation, which includes adhesion receptors, actin regulators, and signaling factors, some of which have roles in mechanosensing (Whittaker et al., 2006; Winograd-Katz et al., 2014; Zaidel-Bar et al., 2007). In this work, we sought to identify and investigate specific adhesome proteins with roles in mechanosensing that potentially drive breast cancer metastasis. Of the candidate genes we screened for, only ezrin, radixin and moesin, the ERM family, were consistently overexpressed in a set of highly metastatic breast cancer cell lines. The ERM proteins are closely related cytoskeletal linker proteins that are pivotal in organizing the interface between the plasma membrane and the actin cytoskeleton, and facilitating processes such as cell adhesion, migration, immune responses, and intracellular signaling (Bretscher et al., 2002; Fehon et al., 2010). The ERM proteins, and particularly ezrin, have been associated with increased metastatic potential in various cancers (Clucas & Valderrama, 2014; Li et al., 2015; McClatchey, 2014; Song et al., 2020). Additionally, the ERM proteins have key regulatory roles in cellular contractility and focal adhesion (FA) maturation (Kunda et al., 2008; Poullet et al., 2001; Quang et al., 2000; Sakurai et al., 2008; Zhang et al., 2020). However, a deep understanding of their functions in mechanosensing, and how this may relate to metastasis progression, is missing. We aimed to shed light on the specific and individual roles of ezrin, radixin and moesin in both mechanosensing and metastasis.

We report antagonistic roles for the ERM proteins in metastasis *in vivo*, wherein knocking down ezrin decreases lung metastasis, while knocking down radixin or moesin increases metastasis. We further demonstrated this antagonistic behavior in various three-dimensional (3D) metastasis assays *in vitro*, using either cell line-derived spheroids or tumor organoids from genetically engineered mouse models (GEMMs) (Padmanaban et al., 2020a). Surprisingly, we found that while ezrin-KD reduces 3D invasion, it also amplifies cellular forces. In contrast, silencing radixin or moesin resulted in increased invasion, as well as decreased cellular forces. We confirmed the ERM-driven inverse relation between cellular force magnitude and invasion using functional experiments. Ezrin-KD was also associated with larger and more adhesions and higher phospho-FAK expression, while radixin-KD and moesin-KD produced opposite outcomes. Additionally, we found that PKC-α,β plays a role in radixin and moesin activation, and inhibiting PKC-α,β was sufficient to restore invasion in the ezrin-KD cells. Taken together, our findings point to a counterbalance between ezrin, radixin and moesin, which regulates adhesions and cellular force magnitude, which in turn acts as a checkpoint for successful invasion and metastasis.

## Results and Discussion

### Ezrin promotes metastasis *in vivo*, while radixin and moesin repress metastasis

Seeking to identify mechanosensing candidates that play a role in breast cancer metastasis, we first used the Human Protein Atlas to identify 32 adhesome genes that were significantly associated with low survival in breast cancer (Wolfenson et al., 2019) (Fig. 1 A). Then, to help identify metastasis-related mechanosensing candidates in breast cancer, we selected 3 pairs of biologically comparable low and high-metastatic breast cancer cell lines: MDA-MB-231 and 231/LM2-4 (Munoz et al., 2006); EMT6 and EMT6-F2 (Shechter et al., 2020); 67NR and 4T1 (Aslakson & Miller, 1992). We carried out a qRT-PCR screen of the 32 genes of interest in these cell lines (Fig. S1 A). Of these candidates, only ezrin, radixin and moesin were consistently significantly upregulated in all three highly metastatic cells, compared to their low-metastatic counterparts (Fig. 1 B), suggesting a role for the ERM gene family in metastasis. Kaplan-Meier plots of patients that exhibited high expressions of ezrin, radixin or moesin reveal relatively low coactivity between the ERM proteins across cancer types (Fig. 1 C), suggesting that their functions may not be fully redundant. This observation challenged the common view of the ERM proteins having largely overlapping roles and encouraged us to investigate their individual roles in breast cancer metastasis. Indeed, whereas ezrin’s function in cancer has been extensively studied, the roles of radixin and moesin as well as the complementary effects of the ERM proteins are poorly understood.

**Figure 1.**
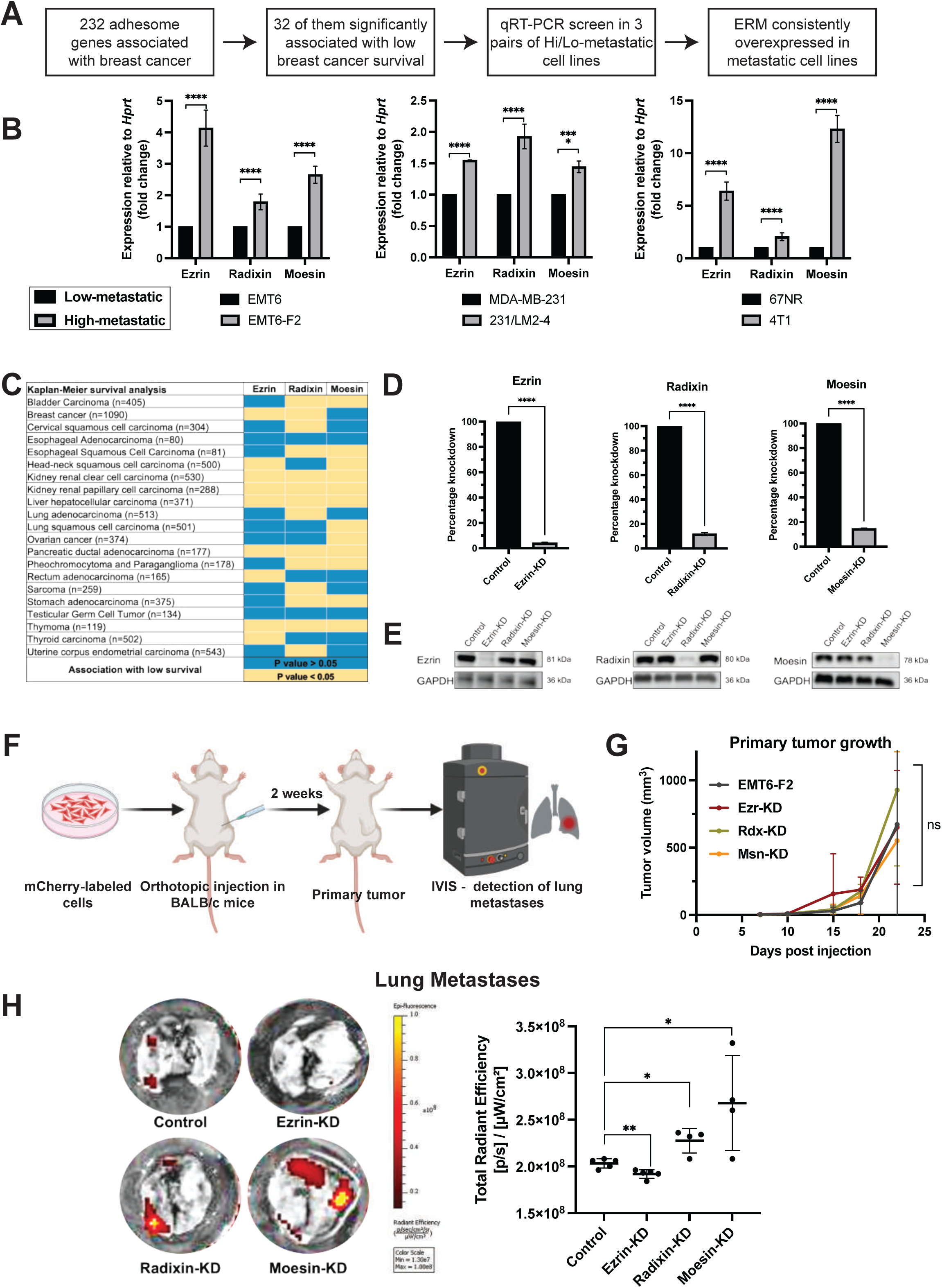
Ezrin promotes metastasis, while radixin and moesin repress metastasis. **(A)** Selection of adhesome genes for a qRT-PCR screen in low versus high-metastatic cell lines: using data from the Human Protein Atlas, 232 adhesome genes were unfavorable with breast cancer (non-prognostic), of which 32 genes were significantly associated with low breast cancer survival(Wolfenson et al., 2019). Findings are summarized in Fig. S1 A. **(B)** qRT-PCR analysis of the expressions of ezrin, radixin and moesin, relative to Hprt expression, and normalized to express fold change. n = 3 biological repeats, ****p < 0.0001, using an unpaired two-tailed t-test. **(C)** The association of a high expression of ezrin, radixin or moesin with survival in 21 cancer types reveals a relatively low prognostic coactivity between ezrin, radixin and moesin. Significantly low survival with P value < 0.05 is marked in yellow and P value > 0.05 is marked in blue. **(D)** qRT-PCR analysis of the expressions of ezrin, radixin and moesin, relative to *Hprt*, showing 85-95% knockdown efficiency compared to control EMT6-F2 cells transduced with non-mammalian shRNA. n = 3 biological repeats, ****p < 0.0001, using an unpaired two-tailed t-test. **(E)** Western blot for ezrin, radixin and moesin expressions in the knockdown cell lysates. At least 80% silencing in each of the knockdowns, n = 3 biological repeats. **(F)** Orthotopic breast cancer mouse model schema: 500,000 mCherry-expressing cells were injected into the mammary fat pad (MFP) of wild type BALB/c mice, then the mice were imaged by IVIS to monitor the development of lung metastases. Created with BioRender.com. **(G)** Tumor growth kinetics graph, as measured by calipers, and calculated using the formula 0.5 x length x width^2^. ns = not significant, p > 0.05 using One-way ANOVA. **(H)** Representative IVIS images of lungs from each group, high signal indicates greater metastatic burden. Graph shows quantification of the metastatic load based on IVIS fluorescent signal (average radiant efficiency). N = 4 or 5 mice per group, *p < 0.05, **p < 0.01 using an unpaired Mann Whitney U test.

Based on their rapid and efficient metastatic spread compared to the EMT6 cells (Fig. S1 B), we selected the EMT6-F2 cells as a basis for our subsequent investigations. To test the individual roles of the ERM proteins, we used lentiviral shRNA transduction to knock down ezrin, radixin or moesin individually in the EMT6-F2 cells. Non-targeted shRNA was used for generating the control cells. Ezrin, radixin and moesin knockdowns were validated by qRT-PCR (Fig. 1 D), Western blot analysis (Fig. 1 E) and immunofluorescent staining (Fig. S1 C).

We next used an orthotopic breast cancer mouse model to test the metastatic potential of the ERM-KD variants *in vivo* (Fig. 1 F). Following injection of mCherry-expressing cells into wild-type BALB/c mice, we monitored the mice for lung metastases using IVIS, and measured the primary tumors twice weekly (Fig. 1 G). The experiment ended 3 weeks later, and the lungs were harvested and analyzed for metastatic foci. Surprisingly, while mice injected with ezrin-KD cells had significantly fewer lung metastases, the radixin-KD and moesin-KD-injected mice had a greater metastatic burden than the control (Fig. 1 H). Thus, these results indicated that radixin and moesin could act as suppressors of metastasis even when expression of the pro-metastatic ezrin was high (Fig. 1 E), highlighting a complex balance between the functions of the ERM proteins.

Importantly, analysis of primary tumor sizes revealed no significant difference between the knockdown groups and the control group (Fig. 1 G). This indicates that the disparity in lung metastasis was not attributable to tumor growth, but rather to other metastasis-promoting factors, such as anchorage-independent cell growth, invasion, or colonization capacities.

### Ezrin promotes invasion and colony formation, while radixin and moesin inhibit both

To elucidate the roles of the ERM proteins in specific metastatic pathways in breast cancer, we established 3D *in vitro* assays that mimic different stages of the metastatic cascade (Padmanaban et al., 2020b). To model cancer invasion into the ECM, which initiates metastasis, we aggregated the cells into 200 µm-diameter spheroids, embedded them into 3D collagen matrices and assessed their invasion capacity 48 h later (Fig. 2 A). In line with the *in vivo* findings, invasion by ezrin-KD spheroids was nearly completely diminished, whereas radixin-KD and moesin-KD spheroids exhibited significantly enhanced invasion relative to controls (Fig. 2, B and C).

**Figure 2:**
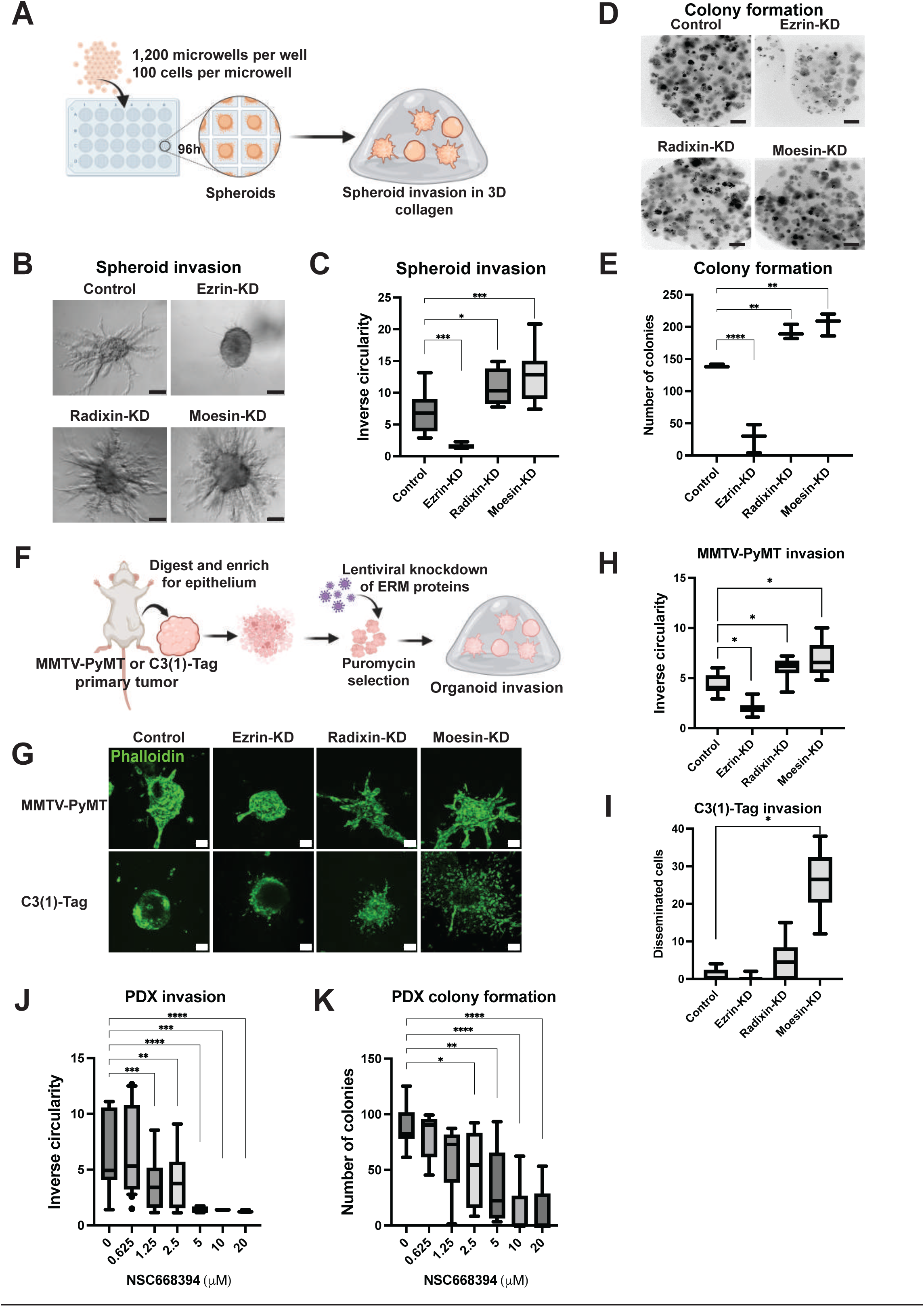
Ezrin promotes invasion and colony formation, while radixin and moesin inhibit both. **(A)** Aggregation of cell lines into spheroids schema: The cells were cultured in suspension in low-adhesion plates for four days, inducing their aggregation into 3D spheroids, each approximately 200 µm in size. The spheroids were embedded in a 3D collagen matrix and cultured for 48 h for invasion. Created with BioRender.com. **(B)** Representative images of a 3D collagen invasion assay of control cells and their ezrin, radixin or moesin-knockdown variants. Spheroids were embedded in collagen, then allowed to invade for 48 h and imaged using a light microscope. Scale bar = 100 µm. **(C)** Quantification of B. The spheroids were fixed, stained with phalloidin, imaged, and their inverse circularity was analyzed to represent invasion. n = 3 biological repeats, *p < 0.05, ***p < 0.001, using One-way ANOVA. **(D)** Single cells were embedded in Matrigel and cultured for one week, or until colonies formed. scale bar = 50 µm. The colonies were fixed, stained with phalloidin and imaged. **(E)** Quantification of the number of colonies. n = 3 biological repeats, **p < 0.01, ****p < 0.0001, using One-way ANOVA. **(F)** Tumor organoid production and transduction: primary tumors were dissected from MMTV-PyMT and C3(1)-Tag mice, then organoids were isolated. Ezrin, radixin and moesin was each knocked down in the organoids by lentiviral transduction. Following puromycin selection (3 days), the organoids were embedded in 3D collagen and allowed to invade. Created with BioRender.com. **(G)** Representative images of invasion of ezrin, radixin or moesin-KD MMTV-PyMT and C3(1)-Tag organoids. Scale bar = 100 µm. **(H)** Quantification of MMTV-PyMT organoid invasion in (G). *p < 0.05, using One-way ANOVA. **(I)** Quantification of C3(1)-Tag organoid invasion in (G). *p < 0.05, using One-way ANOVA. **(J-K)** Quantification of the invasion (J) and colony formation (K) of PDX-derived organoids treated with increasing concentrations of ezrin inhibitor NSC668394. *p < 0.05, **p < 0.01, ***p < 0.001, ***p < 0.0001, using One-way ANOVA.

To model metastatic colonization at distant sites, we next employed a 3D colony formation assay (Padmanaban et al., 2020a). Single cells were embedded in Matrigel and assessed for their ability to generate colonies exceeding 100 μm in diameter. Ezrin silencing significantly decreased the number of colonies formed, whereas radixin or moesin knockdown significantly improved colony formation compared with controls (Fig. 2, D and E). Together, these complementary assays indicate that ezrin and radixin/moesin possess antagonistic functions during the initial and final stages of metastasis.

To validate these findings in clinically relevant spontaneous spontaneous mouse models of breast cancer, or GEMMs (Padmanaban et al., 2020b), we generated tumor organoids from mammary mouse tumors in the luminal MMTV-PyMT model and in the basal C3(1)-Tag model, both of which develop late-stage invasive carcinoma (Guy et al., 1992; Maroulakou et al., 1994). Then, using lentiviral transduction, we knocked down ezrin, radixin, and moesin in the organoids, and performed an invasion assay (Fig. 2 F). The outcome was consistent with the previous results, emphasizing that in both the collective (MMTV-PyMT) and disseminative (C3(1)-Tag) invasion phenotypes displayed by the two models, the ERM proteins significantly and antagonistically determine the extent of organoid invasion (Fig. 2, G-I).

To assess the translational relevance of these findings, we next established organoids from patient-derived xenografts (PDXs) and treated them with NSC668394, an inhibitor selective for ezrin that does not target radixin or moesin. Inhibiting ezrin significantly attenuated both invasion (Fig. 2 I) and colony formation (Fig. 2 J) in PDX organoids, in a dose-dependent manner.

Collectively, these findings establish that ezrin and radixin/moesin exert antagonistic effects on metastatic behaviors across multiple experimental systems, including cell line spheroids, GEMM-derived organoids, and PDX organoids, consistent with our *in vivo* observations. Although NSC668394 was originally identified as an ezrin-binding inhibitor and has shown efficacy without overt acute toxicity in mouse models (Bulut et al., 2011), its mechanisms of action remain incompletely understood. Nevertheless, given the well-established pro-tumorigenic role of ezrin in osteosarcoma, lung cancer, leukemia, and pancreatic cancer (Da Silva et al., 2023), our findings highlight the need to specifically target ezrin in a manner that minimizes inhibition of radixin and moesin, whose functions appear fundamentally distinct and, in this context, opposing.

### The ERM proteins antagonistically regulate rigidity-dependent colony formation

We next set out to delineate the roles of the ERM proteins in specific cellular processes that can drive metastasis. First, we tested their effect on proliferation, and found that silencing the ERM proteins individually had no significant impact on cell proliferation in 2D (Fig. 3 A). We then integrated EdU into the invasion assay to investigate the impact of ezrin, radixin, and moesin expressions on the proliferation rates of cells invading in a 3D context. Similarly, there was no significant difference between the percentage of proliferating control cells and any of the knockdown cells (Fig. 3, B and C).

**Figure 3:**
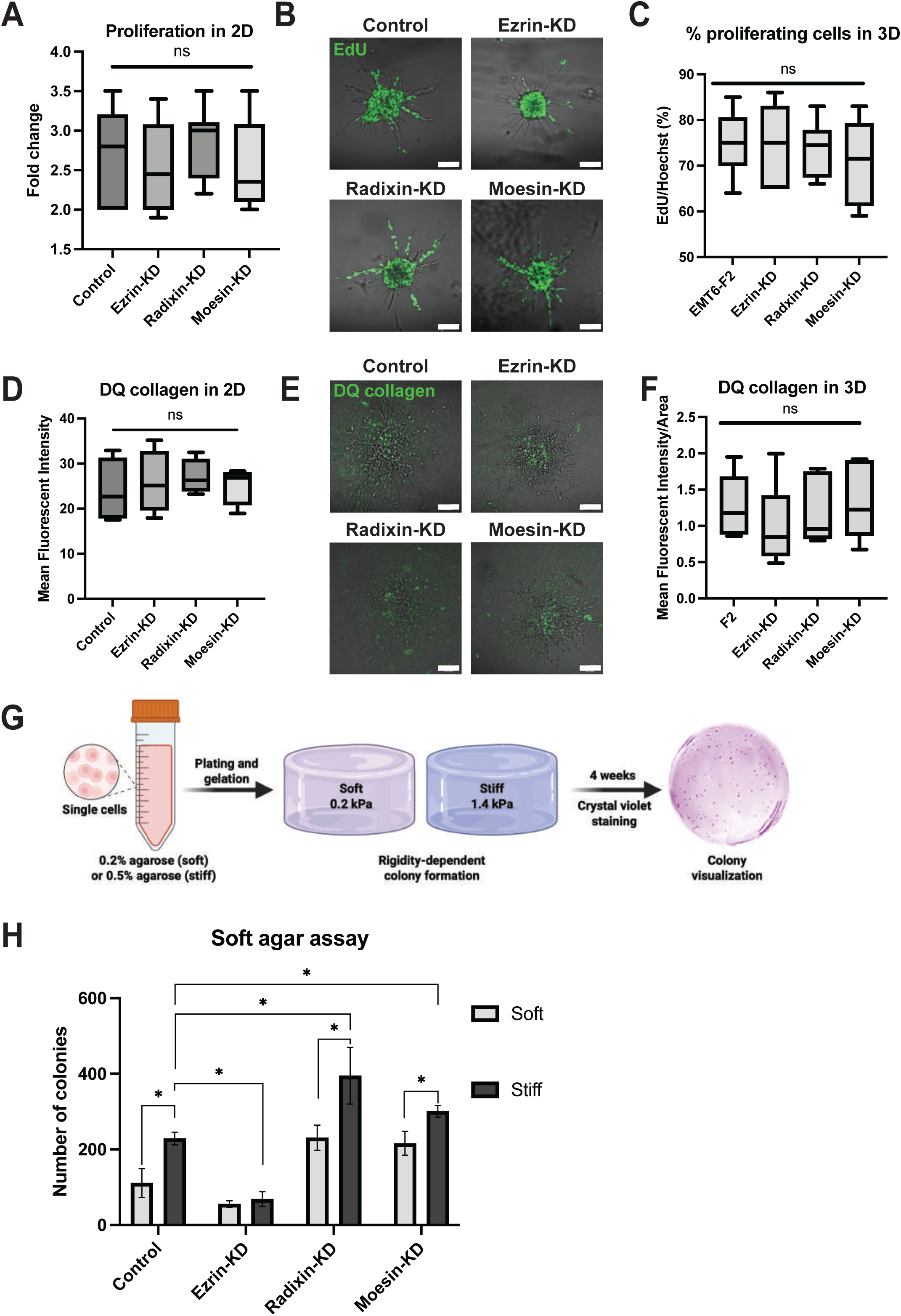
Ezrin promotes rigidity-dependent colony formation. **(A)** Proliferation of cells cultured in monolayer (2D) and quantified as fold change of total cell number at 24 h and 48 h after plating. ns = not significant, p > 0.05 using One-way ANOVA. **(B)** Representative images of EdU-labeled cells in an invasion experiment. EdU-labeled spheroids were fixed 48 h after plating, and stained (green). Spheroids were imaged in z-stack to capture the cells at various depths of the collagen. Scale bar = 100 µm. **(C)** The percentage of proliferating cells was quantified by % EdU-labeled cells out of the total number of cells. ns = not significant, p > 0.05 using One-way ANOVA. **(D)** Quantification of the degree of collagen degradation by cell monolayers, as presented by the mean fluorescent intensity (MFI) within the cell areas. ns = not significant, p > 0.05 using One-way ANOVA. **(E)** Representative images of collagen degradation (green). Spheroids were embedded in collagen mixture with 100 µM DQ collagen and allowed to invade for 48 h. **(F)** Quantification of the degree of collagen degradation by spheroids, as presented by the MFI within the region of the spheroid. ns = not significant, p > 0.05 using One-way ANOVA. **(G)** The control and ERM knockdown cell lines were suspended in 2 percentages of agarose: soft (0.2%, 0.2 kPa) and stiff (0.5%, 1.4 kPa), then cultured for 4 weeks until colonies formed. The colonies were stained with crystal violet, then imaged with a light microscope. Created with BioRender.com. **(H)** Quantification of the number of colonies formed in (G), n = 3 biological repeats, *p < 0.05, using One-way ANOVA.

As ECM remodeling is a critical driver of invasive and metastatic outcome (Butcher et al., 2009), we next tested whether the ERM proteins could alter the collagen degradation capacity of the cells. To that end, we utilized DQ collagen, which yields fluorescent signal upon its cleavage. We found that cells plated on DQ collagen-coated plates displayed no difference in collagen breakdown capacity as a function of ERM expression (Fig. 3 D). We next incorporated DQ collagen into the 3D collagen matrix, and quantified its cleavage during cell invasion. Here too, we found no difference between the collagen breakdown capacities of the invading control spheroids and the three knockdowns (Fig. 3, E and F). Thus, we surmised that the divergent roles of the ERM proteins during invasion and metastasis cannot be attributed to their impact on proliferation or collagen degradation.

Given that the ERM proteins have different impacts on colony formation (Fig. 2) but not on proliferation, we next hypothesized that their antagonistic characteristics may be connected to their functions in mechanosensing or adhesion regulation. To address this, we modified the soft agar assay to elucidate whether rigidity dependence underlies the difference in colony formation capacities. We embedded single cells in 3D agarose gels, that were either soft (0.2% agarose, 0.2 kPa) or stiff (0.5% agarose, 1.4 kPa), and quantified colonies 4 weeks later (Fig. 3 G). In line with the colony formation in Matrigel, we found that regardless of rigidity, the ezrin-KD cells had negligible colony formation, while the radixin-KD and moesin-KD cells each formed significantly more colonies compared to the control cells (Fig. 3 H). Interestingly, the rigid substrate promoted colony formation in all except the ezrin-KD cells, which did not display rigidity dependence or preference (Fig. 3 H). Thus, we concluded that ezrin promotes rigidity-dependent colony formation, strengthening its presumed role in mechanosensing.

### ERM-dependent cellular force transmission is inversely correlated with cancer cell invasion

Having established that the ERM proteins have a role in rigidity-dependent colony formation, we were motivated to explore the relationship between the ERM proteins and cellular force transmission. To that end, we used the PDMS micropillar array system, in which cells were plated onto flexible, fibronectin-coated micropillars. Using live cell imaging, we captured micropillar deflections as the cells adhered and spread upon them. Cellular forces were then calculated using the micropillar displacement measurements (Amer & Wolfenson, 2023; Feld et al., 2020a) (Fig. 4 A). Surprisingly, the non-metastatic ezrin-KD cells generated significantly larger forces compared to the control cells. In comparison, the more metastatic and invasive radixin-KD and moesin-KD cells produced lower forces than the control cells (Fig. 4 B). To functionally test the negative association between invasion and force magnitude, we implemented force perturbation treatments in the course of spheroid invasion (Fig. 4 C). A low dose of blebbistatin (1 µM), an inhibitor of myosin II contractility, significantly increased the degree of invasion in all cell types (Fig. 4 D). Conversely, boosting cellular forces in the spheroids using a Rho activator (CN03A, 1 µg/mL) inhibited invasion (Fig. 4 E).

**Figure 4:**
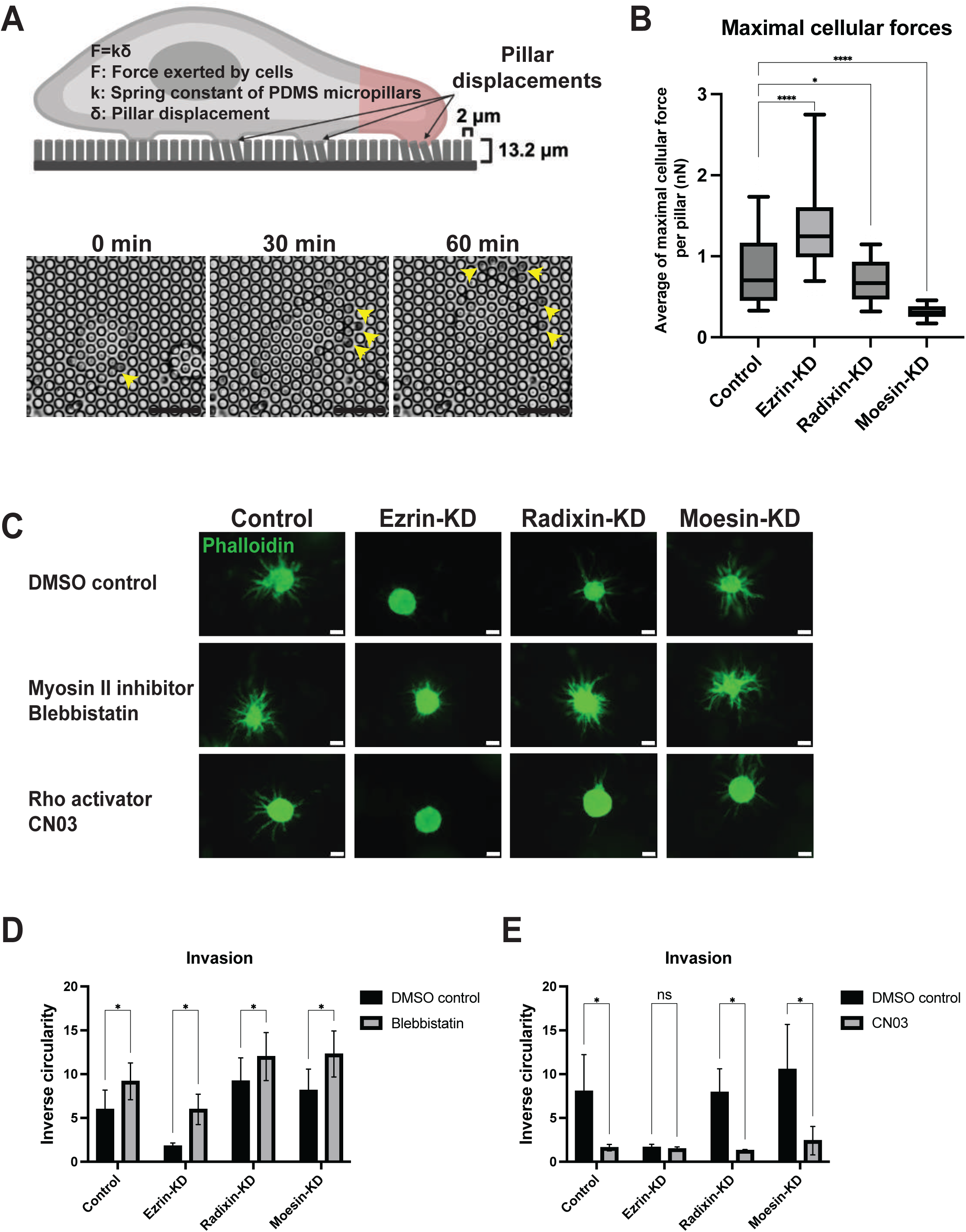
ERM-dependent cellular force transmission is inversely correlated with cancer invasion. **(A)** Calculating cellular forces: cells are plated on PDMS micropillars, then pillar displacements are captured using live cell imaging. Cellular forces are calculated using the formula *F* = *kδ*, where *F* is the force exerted by the cells, *k* is the spring contstant of the PDMS micropillars, and *δ* is the micropillar displacement. *k* is given by the formula 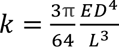, where *D* is the micropillar diameter (2 µm), *L* is the micropillar length (13.2 µm), and *E* is the Young’s modulus of the material (2 MPa for the PDMS substrate)(Feld et al., 2020b). Created with BioRender.com. Lower panel: representative images from live imaging. Yellow arrows indicate micropillar deflections at the start of imaging, at 30 min, and at 60 min. **(B)** Cells were plated on 13.2 µm PDMS micropillar arrays and allowed to settle for 30 min. The cells were imaged using a live microscope for 2-3 h, and the pillar deflections were analyzed. N > 100 pillars per cell, 5-10 cells of each type, *p < 0.05, ****p < 0.0001, using One-way ANOVA. **(C)** Representative images of cellular force perturbation treatments in an invasion assay. Spheroids were fixed 48 h after embedding, stained with phalloidin-488 (green) and imaged. Scale bar = 100 µm. **(D)** Quantification of invasion with 1 µM blebbistatin added to inhibit cellular forces. n = 3 biological repeats, *p < 0.05, using an unpaired two-tailed t-test. **(E)** Quantification of invasion with 1 µg/mL Rho A activator added to increase cellular forces generated. n = 3 biological repeats, *p < 0.05, using an unpaired two-tailed t-test.

Taken together, our results indicate that the invasive and metastatic phenotypes regulated by the ERM proteins were inversely correlated with cellular force generation, in contrast to previous findings (Northey et al., 2017). These findings suggest that excessive force transmission may restrain invasion rather than promote it, and that, rather than maximizing contractility, the ERM proteins regulate metastasis by tuning cellular forces to a permissive range for invasion based on the balance between their coordinated functions.

### ERM proteins regulate adhesions and invasion via FAK and PKC-α,β

Given the well-established coupling between cellular force magnitude and FA growth (Besser & Safran, 2006; Riveline et al., 2001; Stricker et al., 2011), we sought to determine whether ERM knockdowns alter FA characteristics. As the ERM proteins play an important role in adhesion development, which typically occurs at the cell edge, we first examined the membrane localization of each ERM protein under the respective knockdown conditions. As shown above (Fig. 1 E), individual knockdown of the ERM proteins did not alter the overall expression levels of the other two proteins. However, immunofluorescent staining of the different cell populations revealed changes in their membrane localization. Both moesin-KD and radixin-KD cells exhibited increased localization of ezrin at the cell edge, in areas enriched with FAs, and moesin-KD cells also had increased cell edge localized radixin. In contrast, ezrin knockdown resulted in reduced localization of moesin at the edge, with no detectable change in radixin localization (Fig. 5 A; Fig. S3, A and B), suggesting compensatory redistribution among ERM proteins at the cell membrane.

**Figure 5.**
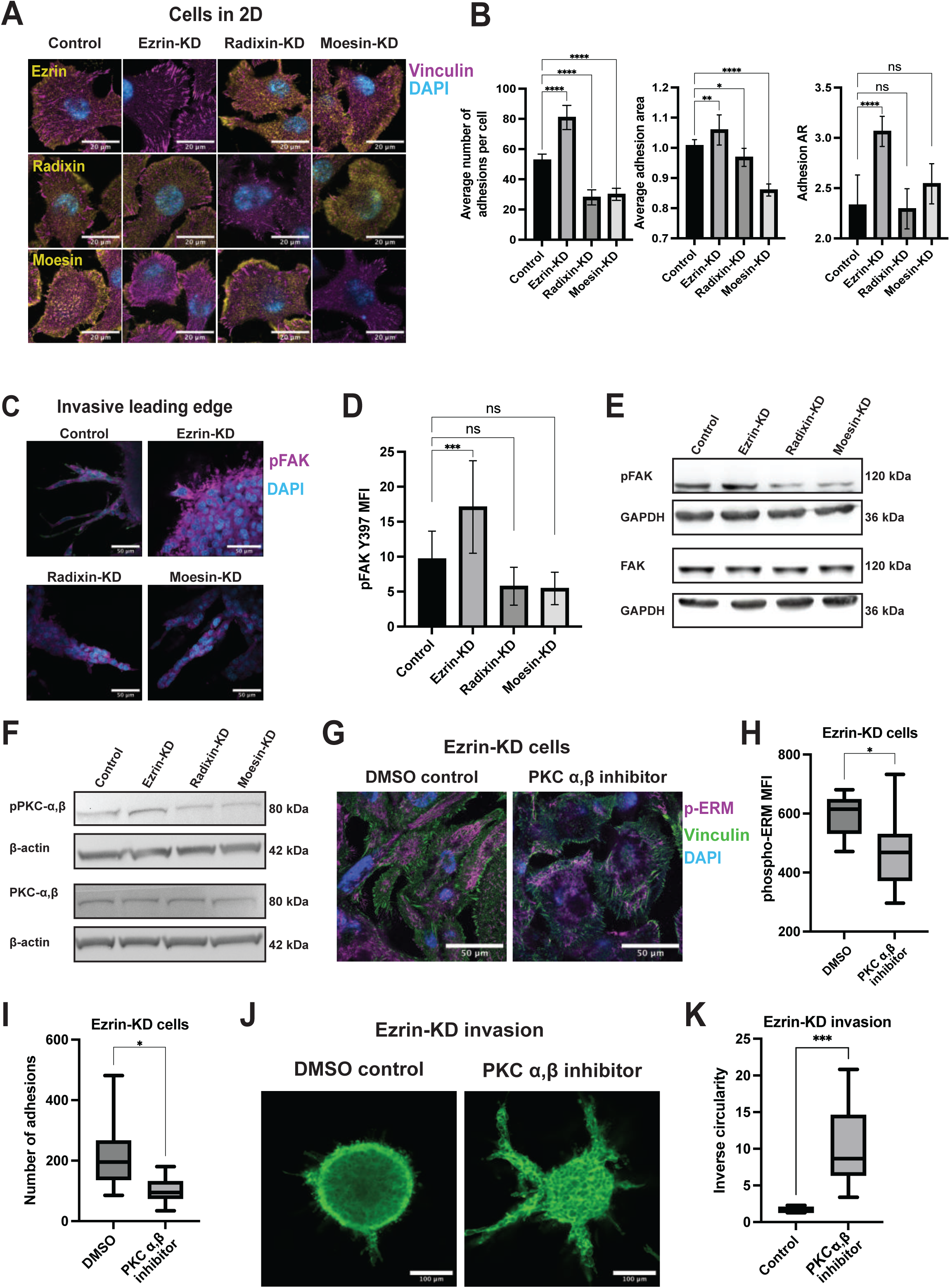
ERM proteins regulate adhesion growth and invasion via FAK and PKC-α,β phosphorylation. **(A)** Representative images of the ERM proteins and FAs in cell monolayers. Cells were plated on ibidi 8-well chambers, then fixed and stained with ezrin, radixin, moesin (yellow) vinculin (magenta), and DAPI (blue). On the right is quantification of ezrin localization at the membrane following individual knockdown of each of the ERMs. **(B)** Quantification of number of adhesions per cell (left) and adhesion aspect ratio (AR) (right) following individual knockdown of each of the ERMs. **(C)** Spheroids were embedded in collagen mixture, then 20 µl was plated in 8-well ibidi chambers. 48 h later, spheroids were fixed and stained with p-FAK (Y357) (magenta), phalloidin-488 (green) and DAPI (blue). Shallow plating allowed for confocal imaging of the invading strands. **(D)** Quantification of p-FAK signal in (C), adjusted to DAPI signal, to account for depth variance across wells and imaged fields. **(E)** Western blot of p-FAK (Y357) and FAK of spheroid lysates. **(F)** Western blot of p-PKC-α,β and PKC-α,β of spheroid lysates. **(G)** Representative images of ezrin-KD cells treated with DMSO control or PKC-α,β inhibitor (Go6976), and stained with phospho-ERM (magenta), vinculin (green) and DAPI (blue). Scale bar 50 µm. **(H)** Quantification of phospho-ERM expression in (G). *p < 0.05 using an unpaired two-tailed t-test. **(I)** Quantification of adhesion number in (G). *p < 0.05 using an unpaired two-tailed t-test. **(J-K)** An invasion assay was set up, then PKC-α,β inhibitor (Go6976) was added. 48 h later, the spheroids were fixed, stained with phalloidin-488, imaged (**J**) and quantified (**K**). n = 3 biological repeats, ***p < 0.001 using an unpaired two-tailed t-test.

We next assessed FA organization by vinculin staining. Strikingly, ezrin-KD cells exhibited a marked increase in abundance of FAs, which were larger and more elongated than those of the control cells. In contrast, radixin-KD and moesin-KD cells had fewer and less mature adhesions than the control cells (Fig. 5, A and B; and Fig. S3 C). Consistent with enhanced adhesion signaling, ezrin depletion was also accompanied by higher phospho-FAK levels in an immunofluorescent staining of invading strands in 3D collagen (Fig. 5, C and D), which was validated by Western blot analysis (Fig. 5 E). Radixin or moesin depletion led to lower phospho-FAK levels compared to controls in the same assays (Fig. 5, C-E). Together, these results indicate that ezrin restrains FA growth, whereas radixin and moesin promote an adhesive state, revealing an unexpected antagonism among ERM family members in the regulation of adhesion maturation. We therefore conclude that the increased recruitment of ezrin to the cell edge upon depletion of radixin or moesin enhances its pro-metastatic function by reducing FA growth and force transmission, making radixin-KD and moesin-KD cells more motile and invasive.

Next, we aimed to investigate signaling pathways which could distinguish ezrin activation from radixin and moesin activation. We used Western blot to analyze the following common ERM activators and mechanosensing candidates: AKT (S473), MYL12A (s19), STAT3 (Y705), 44/42 MAPK, p70 S6 kinase, Src (Tyr416), p38 MAPK, YAP1 (Y357) and MEK1/2 (Ser217/221) (Fig. S3 D). These proteins were not found to be differentially expressed in the ERM knockdown variants. However, Western blot analysis of phospho-Protein kinase C (PKC)-α,β (Koss et al., 2006; Ren et al., 2008), indicated that it was more highly phosphorylated in the ezrin-KD cells compared to the radixin-KD and moesin-KD cells (Fig. 5 F), implicating it in the activation of radixin and moesin. Furthermore, treatment with a PKC-α,β inhibitor, Go6976, decreased phospho-ERM expression in ezrin-KD cells (Fig. 5, G and H), as well as the number of adhesions (Fig. 5, G and I). Finally, inhibiting PKC-α,β was sufficient to rescue spheroid invasion in the ezrin-KD cells (Fig. 5, J and K). This finding suggests that inhibiting PKC-α,β significantly blocks radixin and moesin activation, which overrides ezrin downregulation, and thus promotes invasion.

Collectively, our results indicate that ERM proteins have opposing roles in regulating cell adhesion and force transmission, arising from their competitive recruitment to the cell edge and differential responses to PKC-α/β signaling. These differences lead ezrin to promote metastasis, whereas radixin and moesin suppress it. Consistent with our previous work showing that adhesion maturation is governed by the interplay between extracellular mechanical cues and intracellular mechanosensing machinery (Amer et al., 2021a; Shi et al., 2024), our findings suggest that altered FA dynamics underlie these opposing functions. Parental EMT6-F2 cells, which express high levels of all three ERM proteins compared to the original EMT6 cells, are highly metastatic. However, depletion of radixin or moesin drives them toward an even more aggressive phenotype by leaving ezrin as the predominant cell-edge ERM protein, thereby restraining FA growth and force transmission. Conversely, ezrin depletion increases FA stability and cellular force transmission, resulting in reduced invasiveness. PKC-α/β signaling primarily activates radixin and moesin but may also suppress invasion through ERM-independent pathways that stabilize FAs and enhance force transmission (Bruce-Staskal & Bouton, 2001; Tu et al., 2001). Further studies are needed to identify the intermediate signaling components that connect ERM proteins to FA dynamics and to establish how this regulatory network controls force transmission and tumor cell invasion.

## Acknowledgments

The authors thank Mahasen Sarji for her help with lentiviral transductions, and Yuanning Guo for his help with micropillar analyses. We thank Prof. Yuval Shaked for providing the cell lines. We acknowledge support from the Israel Science Foundation (1738/17, HW) and the Rappaport Family Institute (HW). We acknowledge support from the USA’s National Cancer Institute P30CA006973 (AJE) and NCI U01CA284090 (AJE) and the Breast Cancer Research Foundation (BCRF-24-048, AJE). MA was supported by the Rappaport Technion Integrated Cancer Center (RTICC). MA and AJE were also supported by the Giovanis Institute for Translational Cell Biology.

## Author contributions

M.A. and H.W. conceived the study. M.A. designed the study, performed experiments and analyzed the data. J.V.M. contributed to methodology. T.S. performed experiments. M.A. wrote the manuscript. H.W. and A.J.E. supervised the study. All authors reviewed the manuscript.

## Materials and Methods

### Cell culture, reagents, and transfections

We received the cell lines (MDA-MB-231, 231/LM2-4, EMT6, EMT6-F2, 67NR, 4T1) as a gift from the lab of Prof. Yuval Shaked, Technion – Israel Institute for Technology. Upon receipt, the cells tested negative for mycoplasma. All cells were cultured at 37°C in a 5% CO_2_ incubator in Dulbecco’s Modified Eagle Medium (DMEM), supplemented with 10% fetal bovine serum (FBS), and 100 IU/mL Penicillin-Streptomycin (pen-strep) (Biological Industries). Cell lines were maintained at early passages (<10).

### qRT-PCR analysis

RNA was extracted using the PureLink RNA Mini Kit (12183018A, Invitrogen). Reverse transcription was performed using qScript (95047-100, Quanta Biosciences) to produce cDNA. Quantitative real-time PCR (qRT-PCR) was conducted using PerfeCTa SYBR Green FastMix, ROX (95073-012-4, Quanta Biosciences). Primers (Sigma) are listed in **Table 1**. In all analyses, expression levels were normalized to *Hprt* and relative expression (2^−ΔΔCt^) was calculated.

**Table 1.**
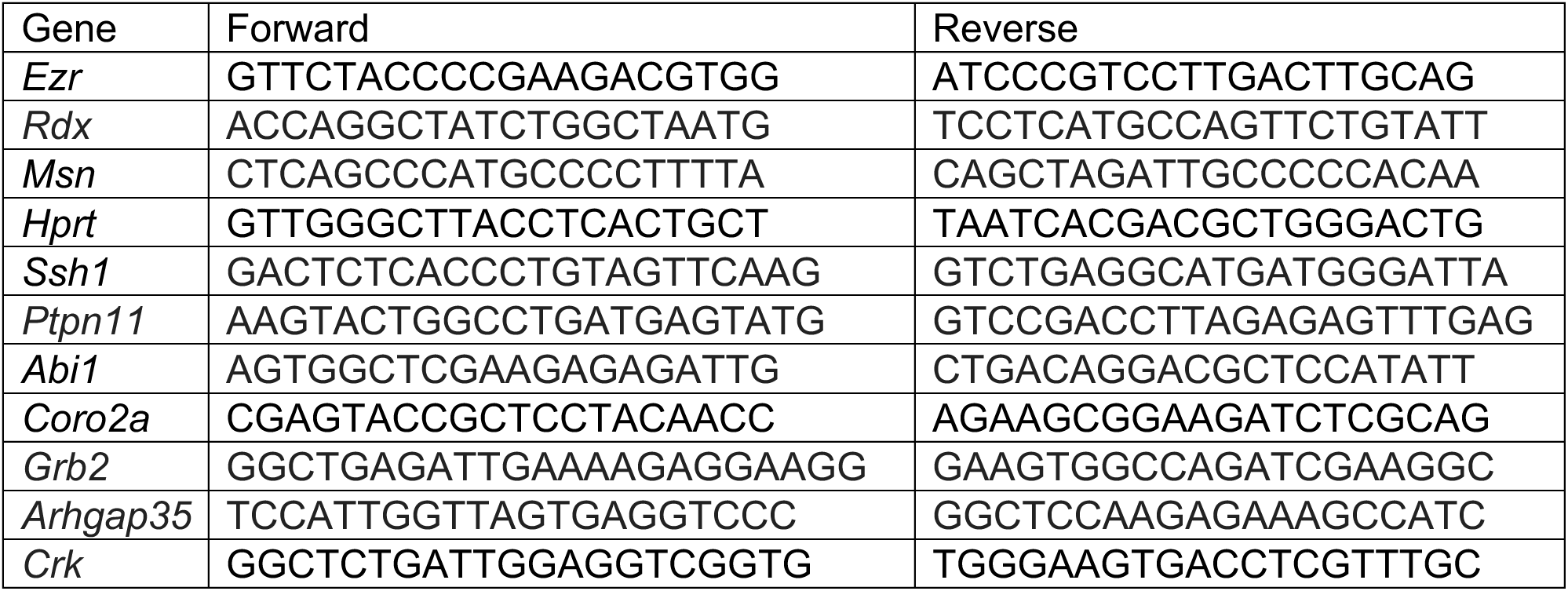
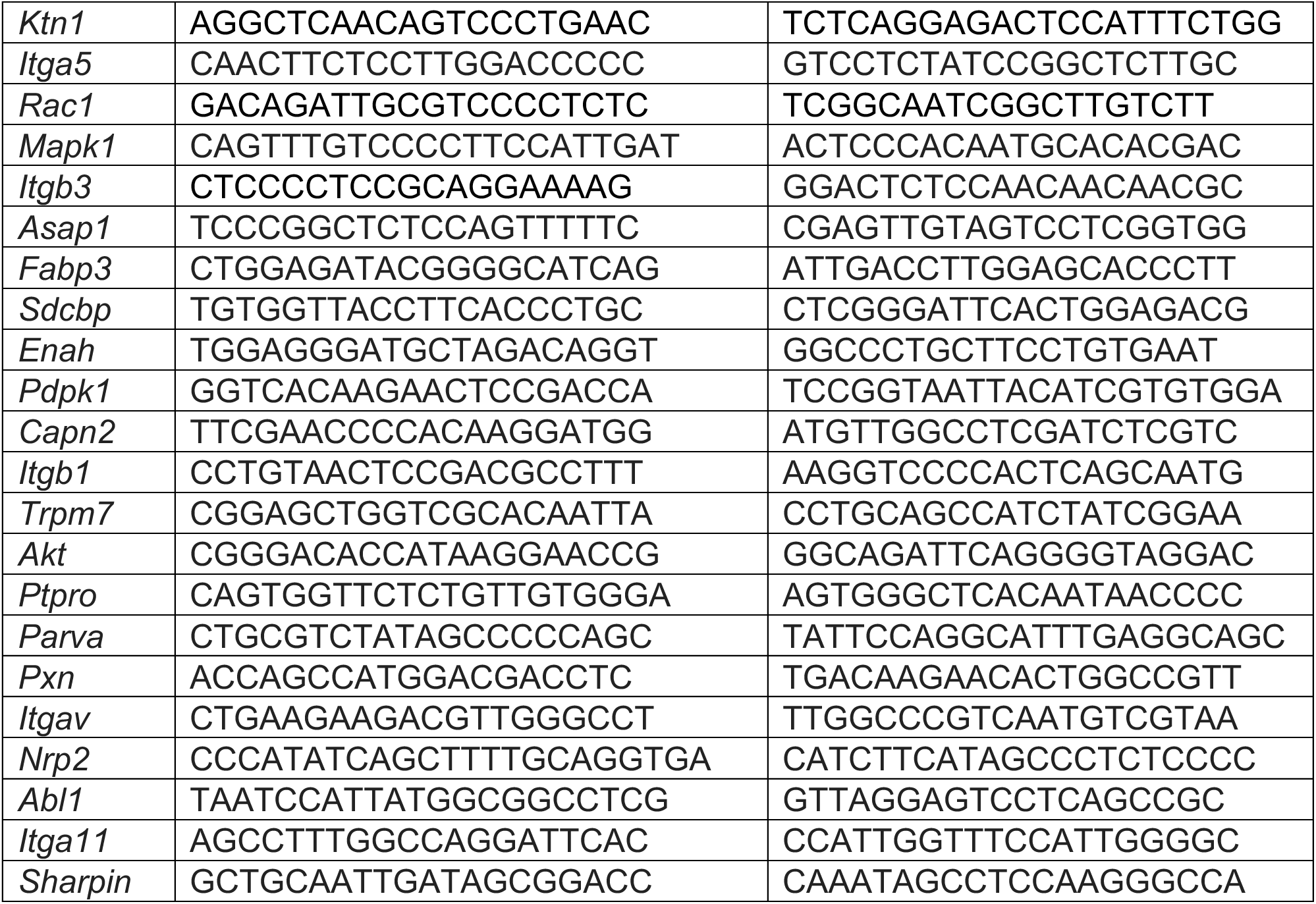
List of qRT-PCR primer sequences used in adhesome screen.

### Lentiviral shRNA transduction

Lentiviral knockdown of ezrin, radixin and moesin was performed using the following shRNA plasmids, from the following bacterial glycerol stocks (Merck, SHCLNG):

- Ezrin: TRCN0000348506, targeted sequence: ACAATGAGCTCAAATCGATTT
- Radixin: TRCN0000352042, targeted sequence: GTTCAGAATTAGCCCAAGCTA
- Moesin: TRCN0000421929, targeted sequence: GAAATCAGGAATATCTCTTTC
- Control cells were generated using the SHC202 - MISSION® TRC2 pLKO.5-puro Non-Mammalian shRNA Control Plasmid (Merck).

First, HEK293T cells were transfected with viral transfer, packaging and envelope plasmids, using the transfection reagent PEI, in Opti-MEM. The media was replaced following overnight incubation, then the cells were cultured for a further 36 h, after which the viral particles-containing HEK293T media was collected and filtered through a 0.45 µm filter. The viral particles were then added to the EMT6-F2 cells for viral infection. After 48 h, puromycin selection (2 μg/mL) was added to the cells. Gene silencing was validated by qRT-PCR, immunofluorescent staining and Western blot.

### Western blot

RIPA Lysis buffer (50 mM Tris-HCl pH 7.4, 1% Nonidet P-40, 0.5% Sodium Deoxycholate, 0.1% SDS, 1 mM EDTA, and 150 mM NaCl) was used to lyse spheroids. Phosphatase inhibitors sodium orthovanadate, sodium fluoride, and protease inhibitor cocktails (#11836153001, Merck) were used. Extracted proteins were loaded into SDS-PAGE gels, and transfer to nitrocellulose membranes was performed using a Trans-Blot SD semi-dry transfer system. The membrane blots were blocked in 5% BSA in TBST for 1 h at room temperature. Blots were incubated with primary antibody overnight at 4°C. After TBST washes, secondary antibodies were added for 1 h at room temperature. Antibodies: Ezrin (Cell Signaling, #3145, 1:1000), Radixin (Cell Signaling, #2636, 1:1000), Moesin (Cell Signaling, #3146, 1:1000), AKT (S473) (Abcam, ab81283, 1:1000), MYL12A (s19) (Abcam, ab2480, 1:500), STAT3 (Y705) (Abcam, ab76315, 1:500), 44/42 MAPK (Cell Signaling, #9101, 1:1000), p70 S6 kinase (Cell Signaling, #9202, 1:1000), Src (Tyr416) (Cell Signaling, #6943, 1:1000), p38 MAPK (Cell Signaling, #9211, 1:1000), YAP1 (Y357) (Abcam, ab62751, 1:500), MEK1/2 (Ser217/221) (Cell Signaling, #9121, 1:1000), GAPDH (ab8245, Abcam, 1:10,000).

### Soft agar assay

Agarose mixtures were prepared at 2x the final percentages shown in **Table 2**, using low-melt agarose (cat # 1613111, Bio-Rad) and deionized sterile water, and then autoclaved. DMEM supplemented with 20% FBS, 2% pen-strep, 2% glutamine was prepared. The agarose mixtures were completely dissolved in the microwave, then kept in a 40°C bath. Each agarose layer consisted of 2X final agarose% + 2X culture media mixed in a 1:1 ratio, for a final volume of 1.5 mL per well. The mixture for the bottom layer was plated in 6-well plates and promptly placed in a cold room for at least 20 min to harden. 5,000 cells were plated in each well: the cells were trypsinized, counted, and resuspended in the agarose mixture for the top layer, then plated on top of the hardened bottom layer, and incubated in a cold room for 20 min. 500 µL cell culture media was added to each well and the plates were incubated at 37°C. The plates were regularly monitored for colony formation, and media was added as needed. Colonies formed within 3-4 weeks of plating. The colonies were fixed and stained using 0.005% Crystal violet + 10% ethanol diluted in PBS. The colonies were imaged under a light microscope and their numbers quantified using the ImageJ (National Institute of Health) cell counter tool. The stiffness of the agarose mixtures was measured by rheometry; 0.2% and 0.5% agarose corresponded to 0.2 kPa and 1.4 kPa respectively.

**Table 2.** Soft agar composition.

| Volume/well = 0.75 mL | Soft agarose (0.2 kPa) | Stiff agarose (1.4 kPa) |
|---|---|---|
| Top layer | 0.2% | 0.5% |
| Bottom layer | 0.5% | 1% |

### Spheroid preparation

Spheroids were prepared using AggreWell™400 plates (Catalog #34450). The wells were coated with Anti-Adherence Rinsing Solution (Catalog #07010) to prevent cell adhesion to the plate, then 1.2×10^5^ cells were plated in each well (100 cells/microwell). The plates were centrifuged to ensure even cell distribution in the microwells, then incubated for 4 days, to allow for spheroid formation.

### Orthotopic breast cancer mouse model

Animals were kept at the Animal Facilities of the Rappaport Faculty of Medicine, Technion (Haifa, Israel), under the strict guidelines of the Institutional Animal Care and Use Committee-approved protocols (IL0860713). Animal studies were performed in compliance with all applicable policies, procedures, and regulatory requirements of the Institutional Animal Care and Use Committee, the Research Animal Resource Center of the Technion, and the National Institutes of Health “Guide for the Care and Use of Laboratory Animals”. The animals were euthanized by CO_2_ inhalation when the experiments were completed. Cells were counted and suspended in PBS at 10^7^ cells/mL, for the injection of 500,000 cells in 50 µL PBS per mouse. 7–8-week-old female BALB/c mice were anesthetized using 4% isoflurane (induction). When the mice were sufficiently anesthetized, the isoflurane level was maintained at 1-2%, and the mice were carefully monitored while under anesthesia. The mice were randomly assigned to the experiment groups and weighed. The cells were mixed by pipetting, then the syringe was prepared for injection. The injection site (4th nipple on the left side) was wiped down with ethanol-soaked gauze, the nipple was clamped using forceps and the needle was inserted bevel-up into the skin. The forceps were released while the needle was stably in position. 50 µL was slowly injected while ensuring that there was no leakage, then the syringe was carefully extracted. The mice were weighed every other day, tumors were measured twice weekly using calipers, and tumor volume was calculated using the formula 0.5 x length x width^2^. The tumor growth and lung metastases were monitored using whole live imaging by IVIS. 3 weeks after injection, when the primary tumors reached a size of 1cm^3^, the experiment was ended, the mice were euthanized, and the lungs were harvested for *ex vivo* IVIS imaging to determine metastatic burden.

### Mice for organoid experiments

For experiments involving PDX, MMTV-PyMT or C3(1)-Tag organoids, all mouse procedures and experiment endpoints were in accordance with protocols approved by the Johns Hopkins Medical Institute Animal Care and Use Committee (ACUC). Mice were euthanized when the tumor burden was less than 20 mm.

### Organoid preparation

The following solutions were prepared:

1. 2.64% BSA/D-PBS
  a. 45.9 mL D-PBS (Sigma # D8662)
  b. 4.1 mL BSA Solution (Sigma #A9576)
  c. Filter sterilize through 50 mL/ 0.22 µm filter flask (Fisher #564-0020)
2. Collagenase solution (30 mL/ tumor)
  a. 27 mL DMEM/ F12 (Life Technologies #10565-018)
  b. 1.5 mL FBS (Sigma #F0926)
  c. 30 µL [50 mg/mL] Gentamicin (Life Technologies #1570-060)
  d. 15 µL [10 mg/mL] Insulin (Sigma #I9278)
  e. 600 µL [0.1 g/mL] Collegnase A (Sigma #C2139)
  f. 600 µL [0.1 g/mL] Trypsin (Life Technologies #27250-018)
  g. Filter sterilize into 50 mL conical tube
3. Organoid media
  a. 490 mL DMEM/F12
  b. 5 mL (100X) Pen/Strep (Sigma #P4333)
  c. 5 mL (100X) ITS (Life Technologies #51500-056)
4. Growth factor media
  a. 10 mL organoid media
  b. 4.3 µL bFGF [2.5 nM] (Sigma #F0291)

The tumor was harvested from the mouse, and the mammary tissue was minced with a #11 scalpel ∼ 50 times, then transferred into the collagenase solution. The collagenase solution was incubated for 1 h at 37°C at 180 RPM in a benchtop shaking incubator. The mixture was centrifuged for 10 min at 1500 RPM, then the pellet was suspended in DNase solution and mixed well. 3 min later, DMEM/F12 was added, then centrifuged for 10 min at 1500 RPM. Differential centrifugation was used to discard single cells, as well as large undigested tissues, leaving only organoids ready for embedding in collagen.

### Invasion assay

The spheroids or organoids were counted and aliquoted in preparation for plating. The collagen mixture was prepared on ice in the following ratio: *375 µL of 10× DMEM + 100 µL of NaOH + 3 mL of 3–3.5 mg/mL rat tail collagen I*.

10 µL collagen mixture was used to coat each well of glass-bottom plates, then incubated for 15 min at 37°C (collagen underlays prevent detachment of the 3D collagen). The plate was placed on a pre-heated plate block set to 37°C, the spheroids were carefully resuspended in the collagen mixture, and 100 µL was plated onto each well. Plates were placed in an incubator for 40 min, to allow the collagen to polymerize, then cell culture media was added to the wells. Inhibitors and treatments were prepared in cell culture media in the concentrations listed in **Table 3**. The plates were cultured for 48 h, then fixed with 4% paraformaldehyde (PFA) and stained with Phalloidin-488. The spheroids were imaged with a Zeiss LSM800 confocal microscope using a 20x 0.9NA, and inverse circularity was quantified using ImageJ.

**Table 3.** Invasion functional assays - inhibitor concentrations.

| Treatment | Working concentration |
|---|---|
| **Ezrin inhibitor** (NSC668394 Calbiochem 341216, Sigma) | 0.625-20 µM |
| **para-nitroblebbistatin (Myosin inhibitor)** (1621326-32-6) | 1 µM |
| **Rho activator** (Cytoskeleton, CN03A) | 1 µg/mL |
| **PKC (α+β) inhibitor** (Go6976) | 10 µM |
| **DMSO control** | Equivalent to DMSO solvent in each treatment |

DQ collagen: DQ™ Collagen, type I From Bovine Skin, Fluorescein Conjugate (Thermo Fisher, D12060) was added to the collagen mixture at 100 µg/mL, according to the manufacturer’s instructions. Collagen digestion by the spheroids during invasion was imaged and quantified.

EdU (proliferation): Click-iT™ Plus EdU Cell Proliferation Kit for Imaging (Alexa Fluor™ 488 dye) (Thermo Fisher C10637) was added to the culture media at 10 µM, according to the manufacturer’s instructions. Proliferating cells which incorporated EdU in their DNA, were imaged and quantified.

### Lentiviral knockdown in organoids

Organoids were suspended in culture media at a concentration of 1000 organoids/100 µl, per well of a 96-well plate. For each transduction, 20 MOI of the viral particles were added to 3 µl viromag (ozbiosciences, VM40100) and the volume was completed to 100 µl per well, then incubated at room temperature for 20 min. The organoids were added to the viral mixture and plated in 96-well non-adherent plates, then incubated on a magnetic plate (OZ Biosciences, MF10000) for 90 min at 37 °C. The plate was then removed from the magnet and returned to the incubator. 24 h later, 150 µL media was replaced with fresh media containing FGF2. The next day, 150 µL media was replaced with media containing 2 μg/mL puromycin. The organoids were cultured in selection media for 3 days, then used for invasion assays.

### Colony formation assay

Corning® Matrigel® Growth Factor Reduced (GFR) Basement Membrane Matrix, LDEV-free, (354230) was thawed overnight at 4°C. Single cells were counted, centrifuged and resuspended in Matrigel, on ice. 200 cells per 20 µL Matrigel were plated in each well of a 96-well glass-bottom plate. The plates were cultured for 1-2 weeks, until colonies formed, then fixed with 4% PFA, 0.25% glutaraldehyde in PBS. The plates were washed, and colonies were stained with Phalloidin-488, then imaged and quantified using ImageJ.

### Pillars fabrication and displacement measurements

The micropillar array molds consist of holes at fixed depths and distances(Ghassemi et al., 2012; Wolfenson et al., 2016b). All pillars had a diameter of 2 µm, and a height of 13.2 µm, and the center-to-center spacing between pillars was 4 μm. Glass-bottom 35 mm dishes (#0 coverslip, Cellvis) underwent plasma treatment. PDMS (Sylgard 184, Dow Corning) mixture was prepared by mixing components A (gel) and B (curing agent) in a 10:1 ratio. A small drop of the PDMS mixture was placed onto each mold, which were then flipped onto the plasma-treated dishes. The dishes were incubated at 65°C for 12 h to cure the PDMS. The molds were then immersed in 100% ethanol, then gently peeled off using a scalpel and forceps. The ethanol was replaced by serial dilutions with PBS: 2 mL ethanol + 7 mL PBS, then 7 mL was aspirated and discarded, and the process was repeated 4 times. Human plasma full-length fibronectin (Merck) was added to the dish at a final concentration of 10 µg/µL, and incubated for 1h at 37°C. Fibronectin was washed away by replacing the buffer with cell culture media, as before. Time-lapse imaging of cells spreading on the pillars was performed using a confocal microscope at 37°C using a 20x objective. Brightfield images were obtained every 30 seconds, for approx. 2 h. Tracking pillar displacements over time was performed with ImageJ using the Nanoracking plugin(Wolfenson et al., 2016c). Pillars that were not in contact with cells at the start of the movie were selected for analysis, and the relative x- and y-positions of each pillar were obtained for all frames. Drift correction was performed using data from pillars that remained undisturbed throughout each movie. The displacement curve of each pillar was generated by Matlab (MathWorks).

### Immunofluorescent staining

Cells were fixed with 4% PFA. Permeabilization was done with 0.5% Triton X-100 in PBS for 10 min at RT, followed by 3x 5 min PBS washes. Blocking was done using 0.1% Tween, 0.2% Triton X-100 in PBS, for 20 min at RT, then washed with PBS. The antibodies (Tables 4, 5) were diluted in blocking solution. The primary antibody solution was added to the cells and incubated overnight at 4°C. Cells were washed three times with PBS, then incubated with the secondary antibody solutions for 1 h at RT. They were then washed with PBS and stained with DAPI (1 µg/mL). Images were taken with a Zeiss LSM800 confocal microscope using a 63x 1.4NA oil objective.

**Table 4.** Immunocytochemistry primary antibodies.

| **Primary antibody** | **Catalog number** | **Dilution** |
|---|---|---|
| Vinculin | Sigma, V9131 | 1:300 |
| Ezrin | Cell Signaling, #3145 | 1:200 |
| Radixin | Cell Signaling, #2636 | 1:200 |
| Moesin | Cell Signaling, #3150 | 1:200 |
| Phospho-Ezrin (Thr567)/Radixin (Thr564)/Moesin (Thr558) (48G2) | Cell Signaling, #3726 | 1:200 |
| Anti-FAK (phospho Y397) antibody [EP2160Y] | Abcam, ab81298 | 1:200 |

**Table 5.**
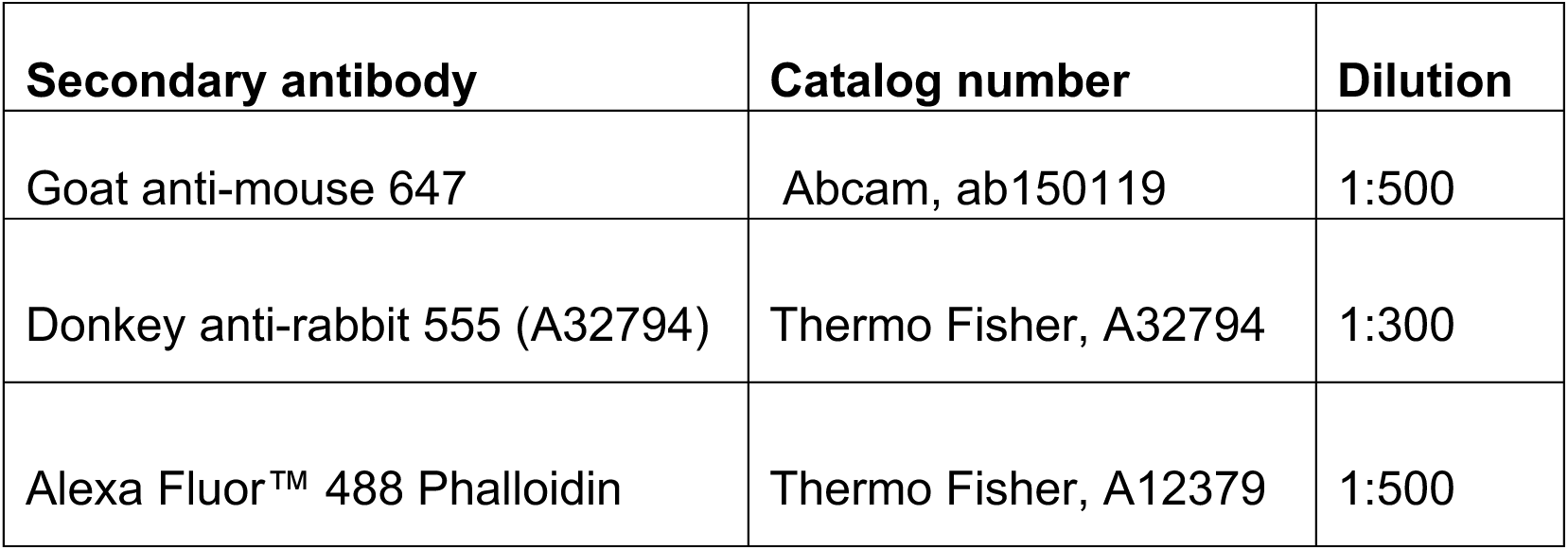
Immunocytochemistry secondary or conjugated antibodies.

| **Secondary antibody** | **Catalog number** | **Dilution** |
|---|---|---|
| Goat anti-mouse 647 | Abcam, ab150119 | 1:500 |
| Donkey anti-rabbit 555 (A32794) | Thermo Fisher, A32794 | 1:300 |
| Alexa Fluor™ 488 Phalloidin | Thermo Fisher, A12379 | 1:500 |

### Statistical analysis

All experiments were repeated three times on separate days using duplicates or triplicates on each day. Significance was calculated using One-way ANOVA or Student’s t-test, as specified in figure legends. Mann-Whitney U test was used to compare IVIS signals in the *in vivo* experiment. Error bars represent standard deviation (sd).

---

**Figure S1.**
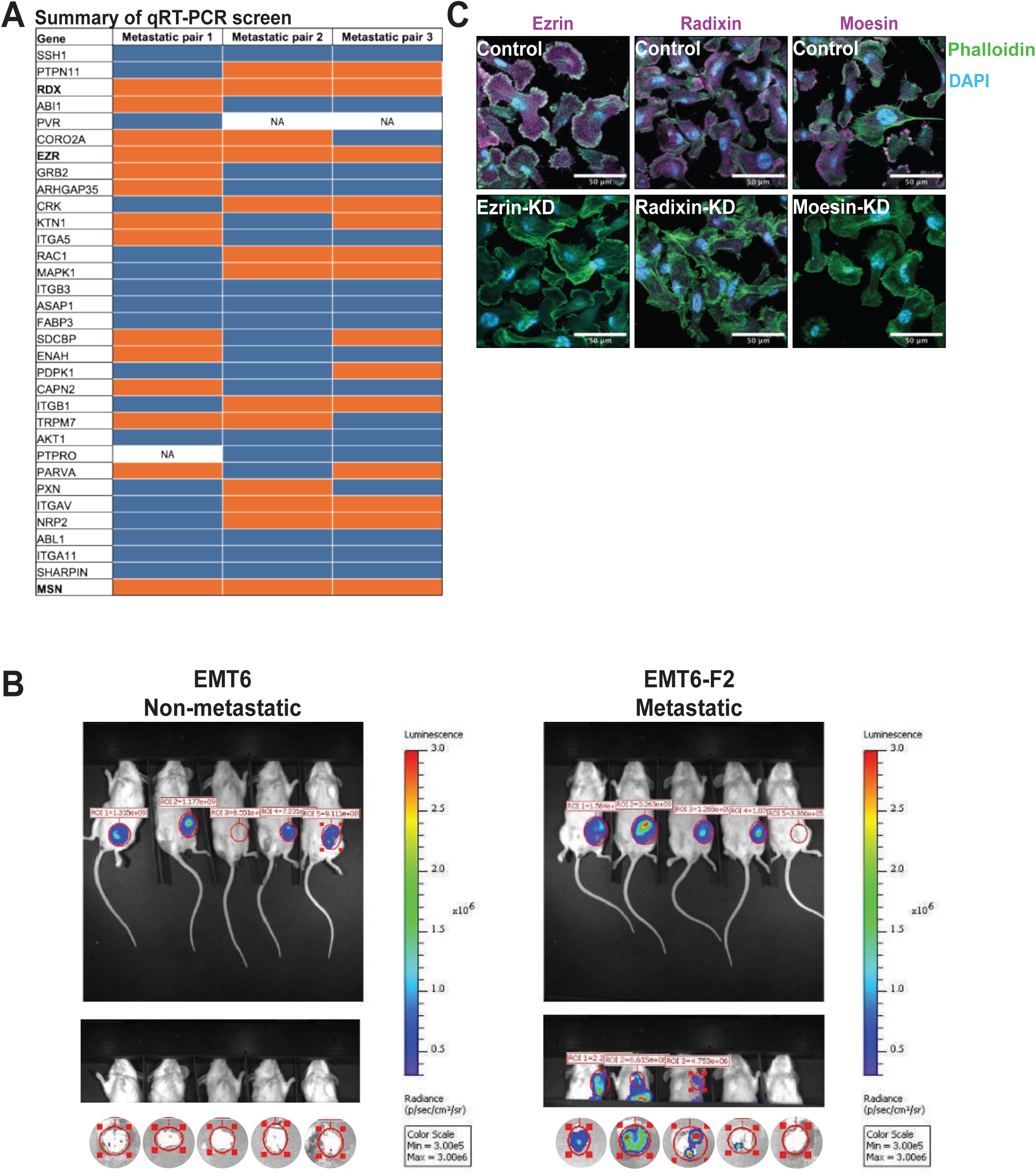
**(A)** Summary of qRT-PCR findings from screen of 32 adhesome candidates in Figure 1A. Metastatic pair 1: MDA-MB-231 and 231/LM2-4; pair 2: EMT6 and EMT6-F2; pair 3: 67NR and 4T1. Orange represents a significant overexpression of the gene in the highly metastatic cell line compared to its non-metastatic counterpart (p < 0.05, using an unpaired two-tailed t-test). Blue represents either a non-significant increase or a decrease in gene expression in the highly metastatic cells. n = 3 biological repeats. **(B)** Validation of the metastatic capacity of EMT6-F2 cell line compared to EMT6 cells. 500,000 luciferase-labeled cells were injected into the MFP of BALB/c mice. The mice were imaged by IVIS to monitor the development of lung metastases. 4 weeks post injection, lungs were harvested and imaged *ex vivo* to acquire signal. In EMT6-injected mice, none of the mice that developed primary tumors had detectable metastases, and in EMT6-F2-injected mice, all the mice that developed primary tumors had detectable lung metastases. **(C)** Representative images of immunofluorescent staining of control, ezrin-KD, radixin-KD and moesin-KD cells, showing a high expression of each of the ERM proteins (magenta) in control cells and low to no expression in the 3 knockdown cell lines. Cells were co-stained with phalloidin-488 (green) and DAPI (blue), scale bar = 20 µm.

**Figure S2.**
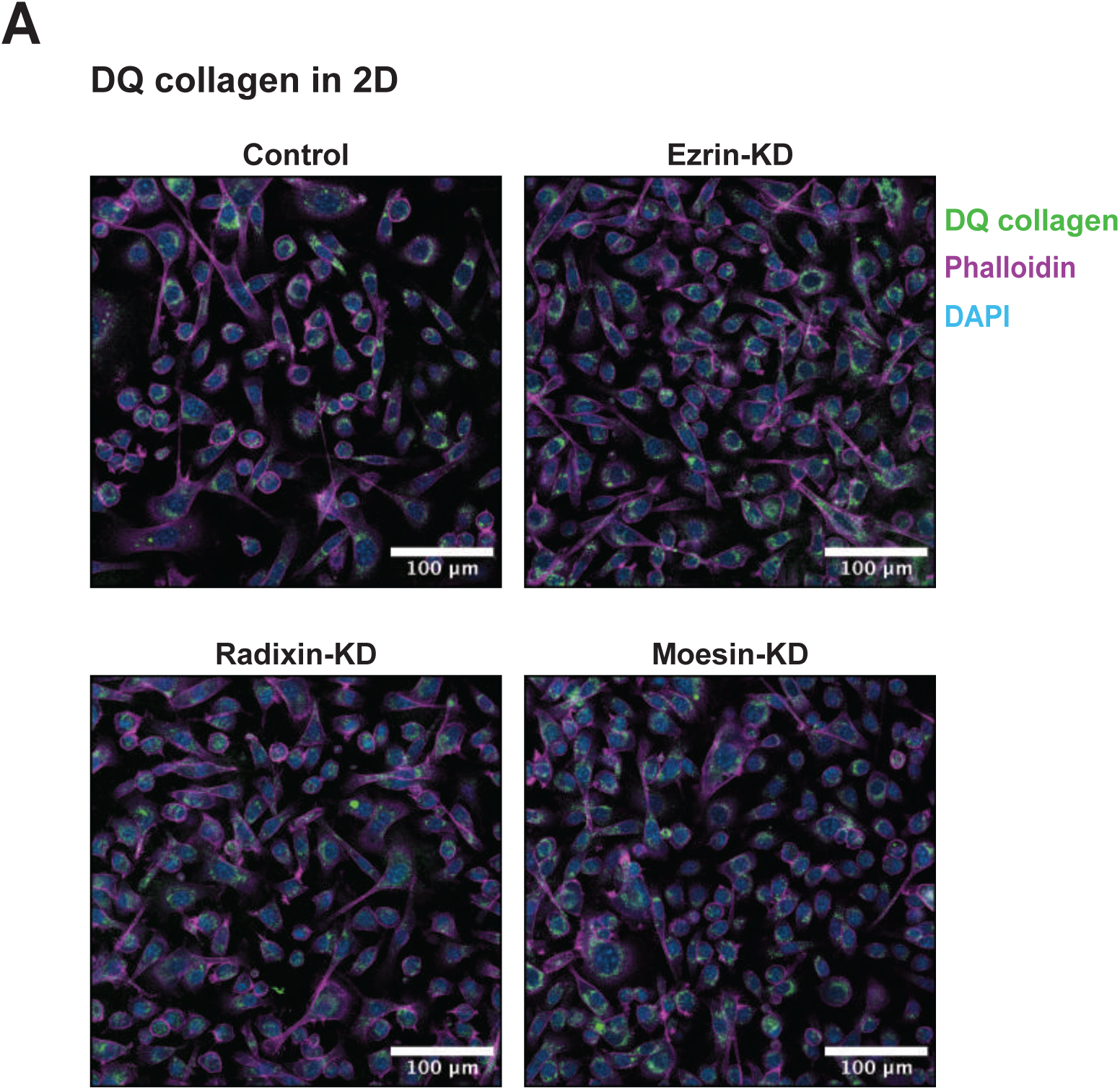
**(A)** Representative images of DQ collagen experiment in 2D (quantification in Fig. 3 D). Wells were coated with a thin layer of collagen-DQ collagen (100 µg/mL) (green) mixture, cells were seeded as a monolayer, then fixed and stained with phalloidin-555 (red) and DAPI (blue).

**Figure S3.**
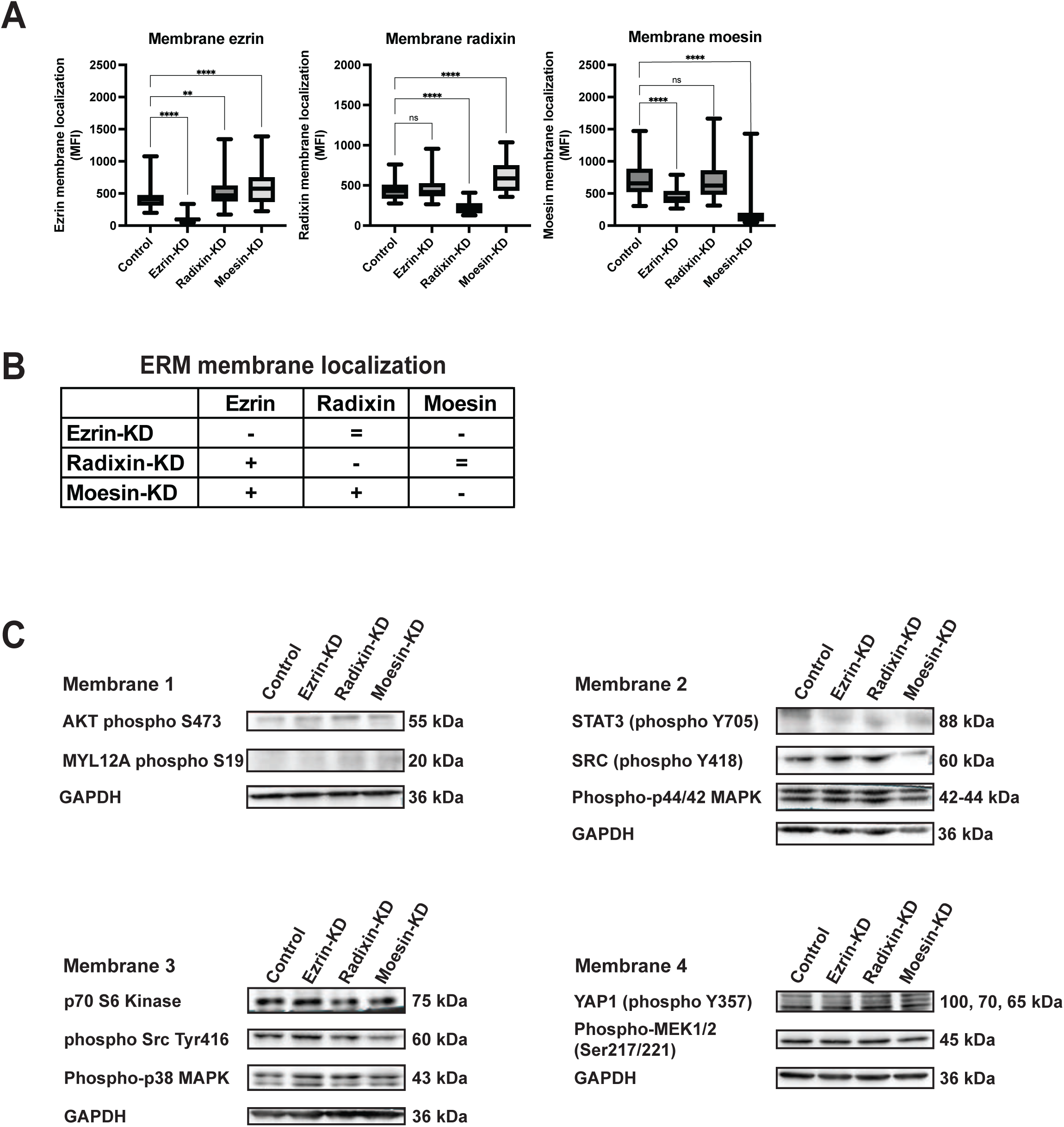
**(A)** Quantification of of ERM localization at the membrane following individual knockdown of each of the ERMs, corresponding to images in Fig. 5 A. **(B)** Summary table of ERM localization at the membrane derived from images in Fig. 5 A and graphs in Fig. S3 A. **(C)** Quantification of adhesion areas following individual knockdown of each of the ERMs. **(D)** Western blots of signaling candidates: AKT (phospho S473), MYL12A (phospho s19), STAT3 (phospho Y705), Src (phospho Y418), phospho 44/42 MAPK, p70 S6 kinase, Src (phospho Tyr416), phospho p38 MAPK, YAP1 (phospho Y357), phospho MEK1/2 (Ser217/221), with GAPDH used as a loading control.

